# Hilbert space filling curves for interpretable point mutation effects on peptide conformational dynamics

**DOI:** 10.1101/2020.01.28.923961

**Authors:** Siddharth S. Rath, Tatum G. Hennig, Tyler D. Jorgenson, Pedro Fischer-Marques, Nitya Krishna Kumar, David Starkebaum, Burak Berk Ustundag, Mehmet Sarikaya

## Abstract

Spontaneous self-organization of solid-binding peptides on single-layer atomic materials offers enormous potential in employing these systems for practical technological and medical applications. Molecular self-organization of peptides depends highly on their sequences which, in turn, affect their conformational behavior under aqueous conditions. Traditional ways of computationally studying the effect of mutations on the conformation states involves dimension reduction on cosine and sine transformed torsion angles, often represented as Ramachandran plots. Although these studies successfully cluster conformation states, they fail to intuitively characterize the effect of the point mutation(s) directly, necessitating further data analysis. Here, we apply Hilbert Space-Filling-Curve (HSFC) on the torsion angles and demonstrate intuitive visualization for the effect of point mutations on conformation states and secondary structure dynamics along a reaction coordinate. We perform molecular dynamics (MD) simulation on wild-type graphene binding peptide (WT-GrBP5). The 12-amino acid long peptide was selected by directed evolution and known to self-organize on atomically flat surface of graphene only under low-neutral pH at room temperature. A charge neutral mutant, M9-GrBP5, on the other hand, assembles at a broader range of pH’s at room temperature, as expected. The HSFC shows clearly that the mutated amino acids in M9 do not correlate with the reaction coordinate of pH change, unlike that of WT, confirming heuristic knowledge. Understanding the effect of specific amino acid φ-ψ pairs that contribute most to the changes in the conformational space of the peptide with changing conditions, will help in analyzing effects of point mutations in peptide sequences. The knowledge of the conformational behavior of solid binding peptides, in general, and its effect on their self-organization propensities on solid surfaces would lead to the rational design of sequences that form soft bio/nano hybrid interfaces in the future towards robust strategies for surface biofunctionalization, in general, and bioelectronics and biosensors, in particular.

## Introduction

Sequence based biomolecules such as proteins, peptides and polynucleotides usually exist in nature with hierarchical structures.[1] The three-dimensional structures adopted by the proteins and peptides, for instance, are intricately associated with the functionality observed in nature and are dependent on the sequence of amino-acids. While proteins show a hierarchy of primary, secondary, tertiary, and quaternary structures, peptides being much smaller, show predominantly secondary structures and are more likely to be intrinsically disordered. [2]

Secondary structure of a peptide is the pattern of hydrogen-bonds present in the structure between the backbone amine and carboxyl groups where hydrogen-bonds are estimated based on Define Secondary Structure of Proteins (DSSP) algorithm. [3] The DSSP algorithm identifies eight distinct categories of secondary structures in polypeptides. Dihedral torsion angles between pairs of amino-acids (AA) in a peptide have been historically used to denote conformations that belong to one of the eight secondary structure clusters. There are three dihedral angles between any two AA in a sequence, namely Φ, Ψ and ω, where Φ and Ψ are free to change between −180 to 180 degrees. ω is usually fixed at 180 or 0 in trans or cis isomers due to the planarity of the peptide bond because of delocalization of lone-pairs on the nitrogen giving it a partial double bond character. [4] Therefore, Φ and Ψ pairs, as defined historically in the context of Ramachandran plots [5] are used to denote backbone conformations ubiquitously. In case of experimental techniques, such as Circular Dichroism (CD), only gross molecular architectures are given such as alpha-helices, beta-sheets, beta-turns, etc., which are more applicable for large proteins as they provide discernible signals.[6] Although signals can be obtained for peptides at higher concentrations, it is still an approximation of the overall conformational ensemble, which is not sufficient to detect the conformation that the peptide attains in the self-assembled structure. For short peptides, that have up to 15-AA long sequences and, therefore, comprise mostly of intrinsically disordered regions, these experimental tools, such as CD, will not provide sufficient information [7]. X-ray crystallography or Nuclear-Magnetic-Resonance (NMR) are difficult and prohibitive to perform, especially for short peptide sequences, and also do not provide high resolution information about the conformation that leads to molecular-recognition.[8] For such short peptides, therefore, and highly specific conformation-dependent self-assembly processes, CD, NMR or X-ray crystallography fall short of the resolution desired.

Several computational studies including Monte Carlo methods [9, 10], Molecular Dynamics (MD) [11–13] and structural alignment for homology modeling [14–16] have been attempted in literature to pinpoint the conformations leading to specific functionality such as those seen in the molecular recognition and self-assembly processes. In each of these methods, secondary structures are almost always denoted in terms of Ramachandran plots of each residue-pair, leading to multiple pairs per sequence. However, such a representation is usually combined with linearization schemes [17–19] and dimension reduction to obtain lower dimensional manifolds [20, 21] of structures. Regular methods of clustering backbone angles, such as principal component analysis [22], multidimensional scaling [23], time-lagged independent components analysis [24] etc., are enough to obtain insight into secondary structures.

Such techniques, however, treat φ and ψ angles for the same pair of residues as separate dimensions and calculate eigenvectors in the linearized cosine and sine transformed torsional angle space. In reality, these angle pairs together characterize the secondary structure propensity of the peptide backbone between the residues and should not be separated when analyzing the effects of individual amino-acids in a sequence. Moreover, the cosine and sine of each backbone angle is not intuitively associated with the effect of point mutations in the sequence on the conformation states and dynamics. Space-filling curves [25–27] are one of the many solutions that can achieve the desired effects of keeping the cosines, sines, and the dihedral angle pairs together, and describe the conformation dynamics in terms of residues directly. In two dimensions, and in a square region, HSFC’s have been shown to be superior to other space filling curves [28–30]. Hence, in the current work, we apply HSFC to the cosine and sine transformed individual φ-ψ angles of a graphene binding peptide WT-GrBP5 and its charge neutral mutant M9 [31, 32]. We select the specific peptides because WT-GrBP5 was selected by directed-evolution [33] to bind to Graphene and shown to assemble on it at room temperature at pH 7. The charge neutral mutant, M9-GrBP5 was rationally designed by replacing Glutamic-Acid (E) and Aspartic-Acid (D) to Glutamine (Q) and Asparagine (N) respectively (Fig 1) to be resistant to electrochemical bias (protonation states) and self-assemble on Graphene at a wide range of pH’s [32].

**Fig 1.**
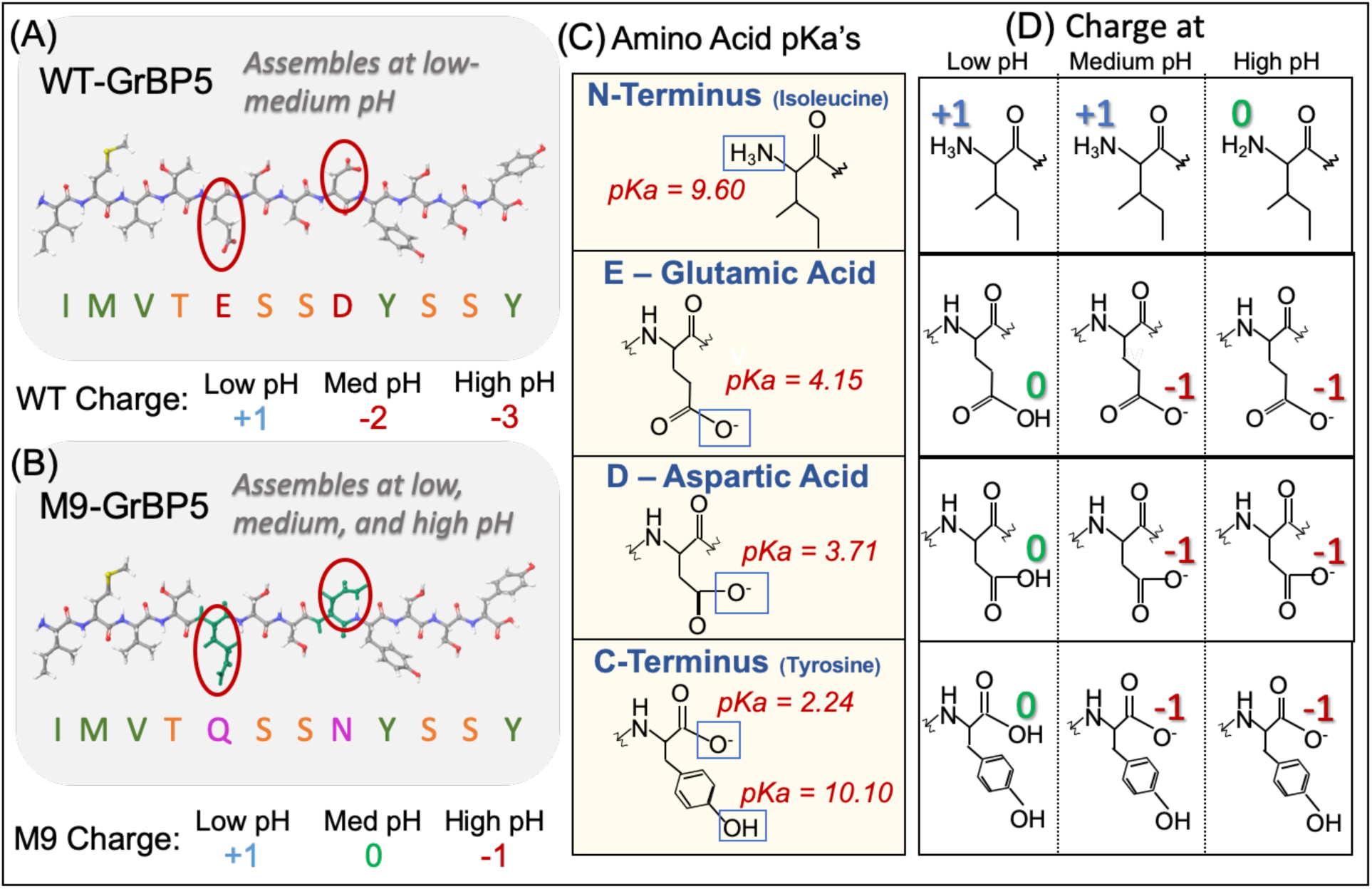
Amino Acid pKa and corresponding protonation states at low, medium, and high pH in WT-GrBP5 and M9 peptides. (A) Sequence of combinatorically selected peptide that self-assembles at room temperature, neutral pH: WT-GrBP5, with charged residues circled. Net charge of the peptide at low pH (pH<2.24), medium pH (4.15<pH<9.6) and high pH (9.6<pH<10.10) are also shown to be +1, −2 and −3 respectively; (B) Sequence of charge neutral mutant of WT: M9-GrBP5, with mutated residues circled. Net charge of the peptide at low pH (pH<2.24), medium pH (2.24<pH<9.6) and high pH (9.6<pH<10.10) are also shown to be +1, 0 and −1 respectively; (C) pKa and protonation sites for N-terminus (Isoleucine), C-terminus (Tyrosine) and charged residues (Aspartic Acid and Glutamic Acid); and (D) Protonation states of relevant amino acids that affect overall net-charge of the peptide, at low, medium and high pH as defined in (A) and (B).

HSFC is applied to keep the backbone dihedral angles (extracted from MD trajectories) together, leading to a decrease in dimensionality four times. The conformation states are now characterized in terms of whole amino-acids instead of cosine and sine of dihedral angles. We do not directly apply a multidimensional HSFC because errors inherent in such a transform increase exponentially with increasing dimensionality [34].

Upon application of the methodology, followed by a principal components analysis (PCA), one obtains an intuitive picture of how the charge neutral point mutations of N and Q instead of E and D do not contribute to conformation changes w.r.t. pH. The current study is focused on intuitively visualizing the effect of the mutations on computationally generated conformation dynamics along the reaction coordinate, which, in this case is pH. We demonstrate that our HSFC approach allows for direct interpretation and visualization of effect of whole amino-acid point mutation on conformation states obtained from molecular dynamics. Such a method will facilitate quick and efficient computational analysis of mutational scans, such as Glycine or Alanine scan of the WT-GrBP5 sequence in the future, to understand the contribution of the amino-acids in the WT-GrBP5 on the self-assembly behavior observed experimentally. Such an understanding of bio/nano interfaces is crucial for designing and understanding bioelectronic devices [35], biosensors and diagnostic tools for a variety of ailments and as such, effective tools to probe dynamic pathways of peptide self-assembly on single layer materials such as Graphene become important.

## Materials and Methods

### Peptides

WT-GrBP5 is a peptide that assembles on Graphene at acidic to neutral pH conditions and room temperature. The sequence is IMVTESSDYSSY. As shown in Fig 1, the three protonation states tested herein include (i) NH_3_^+^– at the N-terminus, –COOH at the C-terminus, protonated Aspartic-Acid (D) and Glutamic-Acid (E), for overall +1 charge (ii) NH_3_^+^– at the N-terminus, –COO^-^ at the C-terminus, de-protonated Aspartic and Glutamic acids, for an overall −2 charge, and (iii) NH_2_– at N-terminus, –COO^-^ at the C-terminus, de-protonated Aspartic and Glutamic acids, for an overall −3 charge. The charge neutral mutant, M9-GrBP5, with sequence IMVTQSSNYSSY and assembly at acidic to basic pH’s, was also modeled at the three protonation states in a similar fashion with overall charges at −1, 0, and +1. Separate molecular dynamics simulations for each pH were then carried out as described below.

The detailed procedures involving experimental peptide containing aqueous solutions, preparation of the graphene substrate, and surface molecular self-assembly have been well described (see references 31, 32, 35, and 41) and will not be repeated herein. Figure 4 schematically describes the peptide assembly along with AFM observation with examples of the self-organized structures of WT and M9 mutant peptides.

### Molecular Dynamics

Molecular Dynamics (MD) simulations of peptides on graphene were carried out using Desmond software [36] in the Biologics Suite of Schrodinger [37] installed on Linux Centos 7.6 and run on an NVIDIA Tesla p100 GPU. The force field OPLS3e is used with all of our simulations and was generated from quantum mechanical calculations [38]. The OPLS3e force field’s small molecule torsion parameter has a coverage of 95% giving it highly accurate predictions of the system’s energetics.

Initial extended peptide structure of WT-GrBP5 was built, protonation state as applicable was assigned, and the force field was applied to the peptide to integrate it into the Schrodinger system by building a disordered system in water. The peptide was modeled at various protonation states in water. Then the most stable solution structure was modeled in the presence of a 15Å x 25Å graphene sheet. pH determination was based on the protonation states of the peptide. The peptide was first modeled in water for 200ns at 300K and medium pH, starting from a minimized extended-structure in a simulation box of 10×10×10 Å^3^ buffer around the peptide. The most-stable structure was changed to reflect protonation states of low and high pH and was remodeled for 200ns in water.

Following the water simulations, simulations with graphene were run for 50ns starting from the most stable water conformation for the particular protonation state, 10 angstroms above the graphene surface with a buffered simulation box of size 1×1×20 Å^3^ around the graphene sheet. All systems were solvated with TIP3P, a 3-site rigid water molecule [39].

We used the NVT ensemble class at 300K for all of our simulations, which fixes the number of particles and keeps volume and temperature constant. For the peptide-water simulations, the relaxation protocol used was constant volume and for the peptide-graphene simulations materials relaxation protocol was used. The system was first minimized using Brownian minimization and then molecular dynamics were performed. Approximately 20,000 frames were collected from each simulation with a time step of 10ps. Trajectory ‘.pdb’ files containing atom coordinates were exported from Schrodinger Biologics suite for analysis. The same procedure was used to set up pH simulations of GrBP5 M9. We acknowledge that the simulation time is not sufficient to exhaustively sample the conformation landscape under the given conditions, however, our purpose is to investigate if the effect of the mutation on the conformational dynamics is still perceptible at such low simulation times. We also observe that the peptide gets kinetically stuck after it adsorbs on Graphene, which may be an artifact of not using a polarizable force-field.

### Principal Component Analysis

The BioPython package [40] in Python was used to obtain the peptide’s backbone torsion-angle pairs, φ and ψ, from the simulation trajectory pdb file. As the peptides are 12-AA long, we obtain 10 pairs of φ and ψ angle pairs per time-frame, excluding the phi and psi angles remaining on each end. Ramachandran plots (Fig 2) of the amino acids in the peptide chain were made using the φ ψ angles of the peptides and compared between the pH simulations to see changes in the backbone conformation (see S1 Fig and S2 Fig for exhaustive Ramachandran plots). To look at the peptide’s backbone conformation as a whole rather than separately between each amino acid, Principal Component Analysis (PCA) was performed on the Linearized φ-ψ space of the peptides.

**Fig 2.**
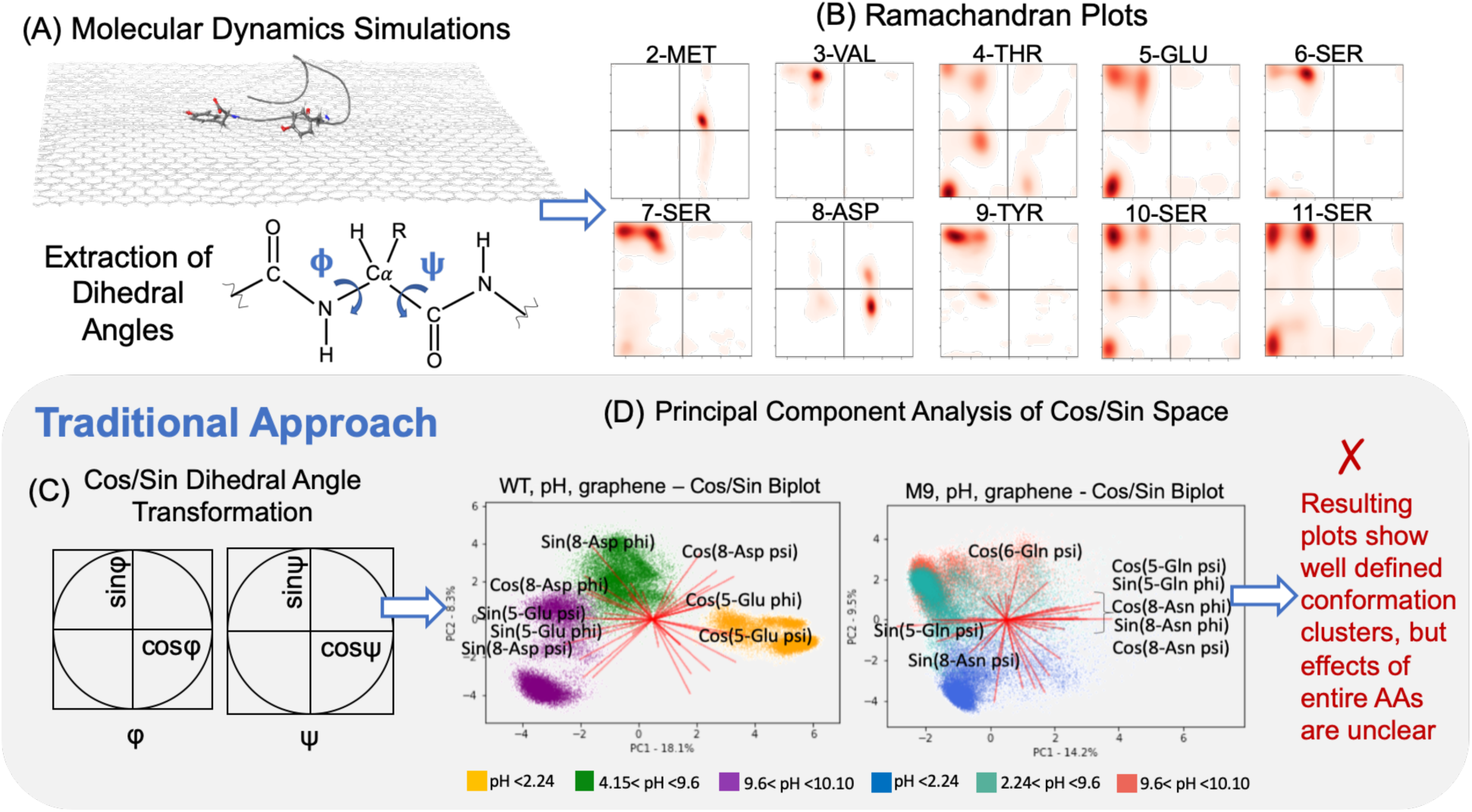
Traditional Molecular Dynamics and PCA Analysis of backbone torsion angles for WT-GrBP5 and M9-GrBP5 at different protonation states. (A) Molecular Dynamics of peptides (WT-GrBP5 and M9-GrBP5 separately) adsorbing on Graphene substrate, NVT ensemble. 200 ns water-structure relaxed for 50 ns on substrate at room temperature. Water model TIP3P. pH is simulated via protonation states as described in Fig 1. Torsion angle pairs for interior amino-acids are extracted (we ignore the psi angle of N-terminus and phi angle at C-terminus); (B) Phi-Psi Ramachandran plots for WT-GrBP5 on Graphene at pH7 to show the multidimensionality of the torsion-angle conformation space of the full peptide; (C) Dimension reduction of full phi-psi space is preceded by linearization via cosine and sine transforms which increases the dimensionality by two times; and (D) PCA Biplots of WT-GrBP5 and M9-GrBP5 on Graphene at different pH (color index in figure) where abscissa and ordinate are the first and second principal components of the cosine and sine transformed phi-psi space. Red lines in the biplots show the original basis vectors projected onto the eigen-basis, with only the charged residues in WT and point-mutations in M9 labeled. They represent the amount each amino acid’s phi/psi angles were factored into the PC1/PC2 eigenvectors. Thus, the longer the line, the larger its contribution to that PC. As we can see, the effect of the charge neutral mutants on the conformation dynamics is difficult to visualize/analyze from the biplots, necessitating further downstream analysis to interpret the individual eigenvectors (principal components).

Linearization is important in order to perform PCA on periodic coordinates. Linearization resulted in overall 40 dimensions in our case, viz., cosine and sine of each φ and ψ angle. PCA is an eigenvector-based multivariate analysis that uses an orthogonal transformation to explain the variance of the given data set. First the cosines and sines of the φ ψ data, from the simulations, was combined into one file. The data was then scaled and centered by changing the average value to zero and the standard deviation to one. The PCA object was then created and the Principal Component (PC) coordinates and their explained variance are calculated (See S3 Fig and S4 Fig). Focusing on PC-1 and PC-2, since those principal components have the highest explained variance, clusters of backbone structures can be revealed in PC-1 vs PC-2 scatterplots. The loading scores along PC-1 and PC-2 were found to determine which φ and ψ angles were contributing the most to the variance among clusters. Biplots (Fig 2) were created to overlay PC-1 and PC-2 space with vectors corresponding to the influences that each dimension has on the principal components. The loading scores for PC-1 and PC-2 are in S5 Fig and S6 Fig for WT-GrBP5 and M9-GrBP5, respectively.

### Hilbert Space-Filling Curves

In order to look at the φ and ψ angles of a single amino acid as a whole rather than separately, the cosines and sines of the φ and ψ angles were transformed to one-dimensional coordinate using an order 10 Hilbert space filling curve successively through the cosφ-sinφ and cosψ-sinψ space (Fig 3). Hilbert curves are useful in transforming between 2D and 1D while preserving the relative locality of a point in the 2D space.

**Fig 3.**
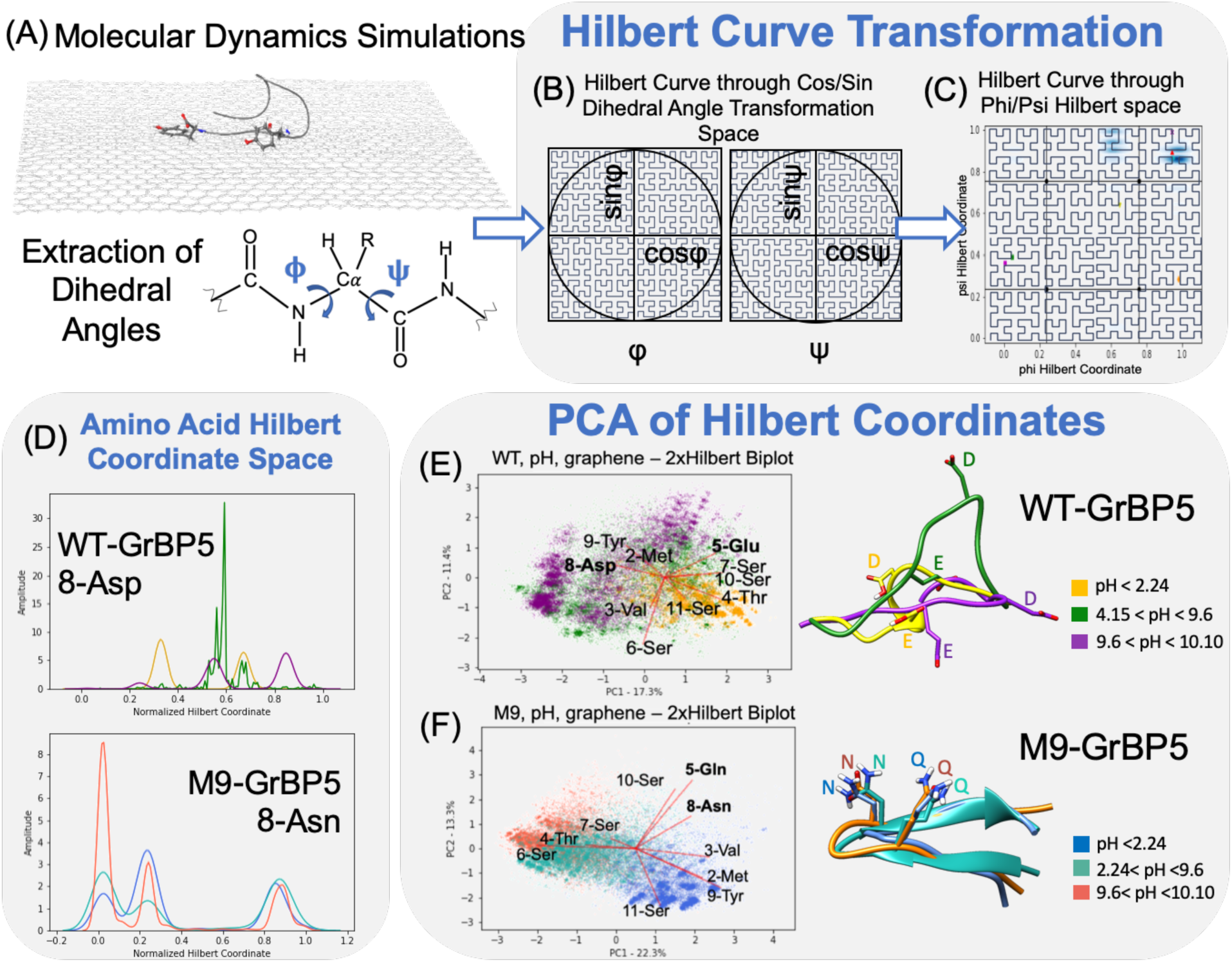
Application of Hilbert Space-Filling Curve (HSFC) on torsion angles followed by PCA for WT-GrBP5 and M9-GrBP5 at different protonation states. (A) Molecular Dynamics of peptides (WT-GrBP5 and M9-GrBP5 separately) adsorbing on Graphene substrate, NVT ensemble. 200 ns water-structure relaxed for 50 ns on substrate at room temperature. Water model TIP3P. pH is simulated via protonation states as described in Fig 1. Torsion angle pairs for interior amino-acids are extracted (we ignore the psi angle of N-terminus and phi angle at C-terminus); (B) Application of HSFC of order 10 on cosine(φ) vs sine(φ) and cosine(ψ) vs sine(ψ) space to consolidate the two components of a torsion angle into one: either phi or psi, respectively; (C) Application of HSFC of order 10 on the consolidated torsion angle space to consolidate φ-ψ pairs around every amino-acid into one variable prior to performing Principal Components Analysis (eigen-basis transformation); (D) Hilbert coordinate (consolidated φ-ψ pair variable) for Aspartic-Acid (Asp) or Asparagine (Asn), where φ is between Asp/Asn and Serine (Ser) and ψ is between Asp/Asn and Tyrosine (Tyr) in the sequence IMVTESSDYSSY (WT-GrBP5) and IMVTQSSNYSSY (M9-GrBP5) respectively. Details and legend are shown in Fig 4 and S7 Fig & S8 Fig; (E) and (F) PCA of φ-ψ pair variables (Hilbert Coordinate, HC) for all the amino-acids in WT-GrBP5 and M9-GrBP5 on Graphene at low, medium, and high pH, respectively. pH is simulated in terms of protonation states. Red-lines show the original pair-variable (HC) dimensions, for each amino-acid as labeled, projected onto the eigen-basis of the first two eigenvectors (principal components). The stable structures obtained from the individual MD trajectories for the different pH’s are shown aligned on each other and color coded as shown.

The Hilbert transformation assumes the space is divided into n by n pixel cells with the coordinate 0 in the lower left corner and n^2^– 1 coordinate in the lower right corner. For an order 10 Hilbert curve, the cosine-sine space for each dihedral angle was converted into a box with 1024×1024 pixels. To transform the data into Hilbert space, the following equation was used:

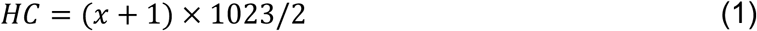

where HC is the Hilbert Coordinate, a paired-variable. This equation shifted the cosine and sine value range of −1 – +1 to 0 – 2 and determined the Hilbert coordinates (Fig 3). This operation also reduced dimensions two times. A second Hilbert transform was applied on the transformed data to reduce dimensionality two times more and PCA was then performed on the data to create biplots.

## Results and Discussions

The variability in torsional angles of D and E in WT-GrBP5 is clearly captured in the structures obtained and from the biplots shown, it can be seen that both D and E are changing along the reaction coordinate which is the increasing pH direction. Similarly, the invariance of torsion angles of charge neutral mutants Q and N in M9 that replace E and D in WT with pH is clearly seen in the structures as well as in the orthogonality of 5-Gln and 8-Asn vectors to the reaction coordinate in the biplot in Fig 3F. The biplots in Fig 3 clearly show that the individual charge neutral point mutations are not affected by protonation changes and are not contributing to conformation dynamics along the pH reaction coordinate, as expected from prior experimental results from previous work done [32].

The Ramachandran plots of Aspartic Acid and Asparagine in Wt-GrBP5 and M9-GrBP5 respectively, are shown in greater detail in Fig 4; along with the corresponding HSFC plots, to emphasize the advantages of visualizing φ-ψ pair dynamics in a consolidated manner.

**Fig 4.**
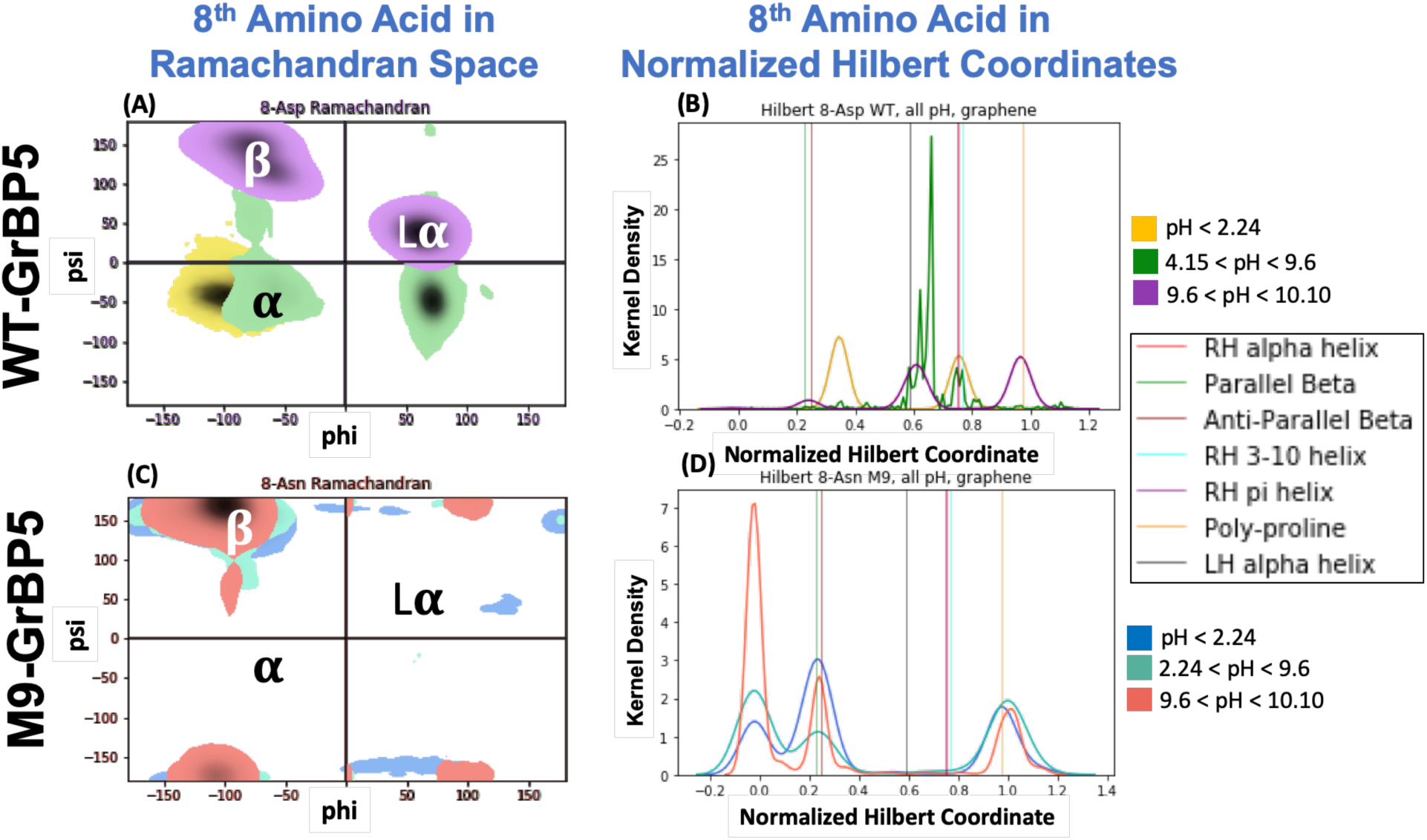
Ramachandran Plot vs Hilbert Space-Filling Curve (HSFC) on torsion angles prior to PCA Analysis for WT-GrBP5 and M9-GrBP5 at different protonation states. (A) Ramachandran plot for φ-ψ distributions for low pH (yellow), medium pH (green) and high pH (purple) for WT-GrBP5. Intensity of the colors show the density of the φ-ψ pair; (B) Normalized HSFC plot for φ-ψ distributions for low pH (yellow), medium pH (green) and high pH (purple) for WT-GrBP5. Since φ-ψ pair is now consolidated into the x-axis, the y-axis is the density, or prevalence of the φ-ψ pair. The vertical lines identify specific, well-defined secondary structures, which are easier to pinpoint in the HSFC plots (legend shown); (C) Ramachandran plot for φ-ψ distributions for low pH (blue), medium pH (cyan) and high pH (red) for M9-GrBP5. Intensity of the colors show the density of the φ-ψ pair; (E) Normalized HSFC plot for φ-ψ distributions for low pH (blue), medium pH (cyan) and high pH (red) for M9-GrBP5. The vertical lines identify specific, well-defined secondary structures, which are easier to pinpoint in the HSFC plots (legend shown).

The molecular self-organization of the peptides take place under ambient aqueous conditions when they are placed on freshly cleaved atomically flat graphite sample, as schematically demonstrated in Fig 5, a standard process that was discussed extensively in prior publications.[31,32,41] Also shown in this figure are the experimentally acquired images using atomic force microscopy (AFM) on samples that were incubated 1μM concentration of WT-GrBP5 and M9-GrBP5 separately on HOPG for 1 hour at pH 3.5, pH 7 and pH 9.7, with buffers H_3_PO_4_, NaH_2_PO_4_, and NaOH respectively.

**Fig 5.**
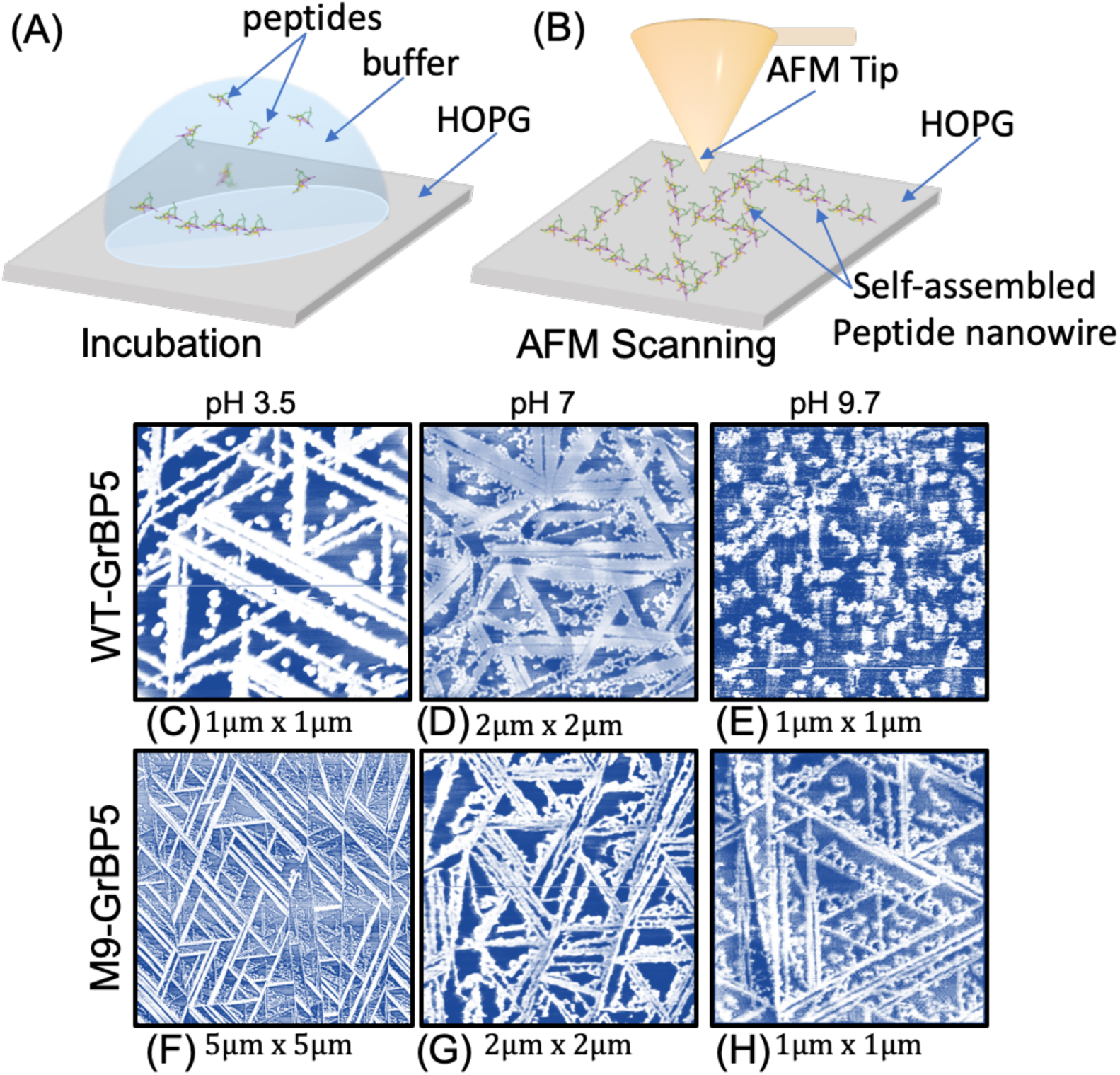
Experimental evidence of occurrence of Self-Assembly in WT and M9 GrBP5 peptides on Graphite (HOPG) surface under different pH. (A) Incubation of 1uM concentration of peptides dissolved in buffer (NaH_2_PO_4_ for pH 3.5, DI water for pH 7 and NaOH for pH 9.7) on HOPG surface for 1hr each; (B) Atomic Force Microscopy scanning of the surface after incubation and lyophilization, to obtain topography as a result of electrostatic interactions between tip and surface; (C) topography of WT-GrBP5 on HOPG sample at pH 3.5, room T showing the presence of long range ordered structures; (D) topography of WT-GrBP5 on HOPG sample at pH 7, room T showing the presence of long range ordered structures; (E) topography of WT-GrBP5 on HOPG sample at pH 9.7, room T showing the presence of islands and clusters but no long-range ordered structures; (F)-(H) topography of M9-GrBP5 on HOPG sample at pH 3.5, 7 and 9.7 respectively at room T, showing the presence of long range ordered structures. White regions are peptide while the darker regions are the substrate in all the AFM scans. All scale bars are 200nm in length and all scan areas shown are 1μm^2^.

The presence of self-assembled peptide nanostructure is observed, i.e., long range ordered domains, on HOPG for WT-GrBP5 at low and neutral pH, but the phenomenon is not observed at higher pH’s. In contrast, the rationally designed charge-neutral mutant displays spontaneous self-assembly at all the pH’s tested. Such effects on functionality are usually dictated by conformation ensembles which are in turn affected by mutations in the sequence. Our HSFC method can help in quickly identifying whether or not a point mutation is strongly correlated with the conformation dynamics along the desired reaction coordinate, thereby enabling a platform to computationally perform alanine scans, for example, with intuitive interpretations and visualizations of the MD trajectories.

## Conclusions and Future Work

In the current work, we apply Hilbert Space Filling Curves (HSFC) on torsion angles from molecular dynamics trajectories and demonstrate its capability in analyzing the effects of point mutations on conformations. Applying the HSFC approach directly to the φ-ψ Ramachandran space does not result in a meaningful separation of conformations, and just projecting φ-ψ into cosφ-sinφ and cosψ-sinψ space is enough, especially when overarching differences matter more than small differences, as in the case of folding-unfolding pathways. However, more intuitive information about effects of point mutations can be acquired from the same simulations, upon keeping the φ and ψ together for each pair of residues in the sequence.

Identification of whether a point mutation is strongly correlated with the conformation dynamics along the desired reaction coordinate are crucial for analyzing amino-acid scans computationally. Rapid and intuitive analysis of effects of point-mutations on conformation states, as in an Alanine or Glycine scan, are crucial in understanding interfacial interactions that guide self-assembly of peptides on atomically flat solid materials. Despite the lack of long simulation times, the approach was still capable to produce results, in just 50 ns of graphene-peptide simulations. We still observe that the backbone angles of the charge-neutral mutants were not correlated with the reaction coordinate of pH or protonation state. The conclusion is consistent with the reasoning behind the rational design of the self-assembling M9-GrBP5 mutant.

In the future, the methodology introduced herein could be performed on Metadynamics simulation data of adsorption of WT-GrBP5 on Graphene. The simulation will be performed with a polarizable force field. Alanine or Glycine scan mutants of WT-GrBP5 will also be simulated as such, to probe the effect of amino-acids that are most correlated with conformation dynamics as the peptides self-assemble on Graphene and correlate them to experimentally observed trends. These forthcoming studies could provide the fundamentals for computational design of novel peptide sequences as possible candidates to form spontaneous long-range ordered self-assembly on graphene and other single layer materials where the molecular assembly play the major role in peptide-enabled devices of the future, In particular, the knowledge of precise soft bio/nano interface structure could aid *ab-initio* calculations in the future to explain the effect of self-assembly on the electronic property changes in the bio/nano hybrid systems to design biosensors, bioelectronics, bio-photovoltaics, and biomolecular fuel cells.

## Code and Data Availability

The PCA and HSFC code used in the study are available on GitHub at https://github.com/Sarikaya-Lab-GEMSEC/HSFC_MD

## Acknowledgments

The research was supported by the DMREF Program at National Science Foundation (NSF) through the MGI platform (Materials Genome Initiative) under grant numbers DMR# 1629071, 1848911, and 1922020.

## Supplementary Information

**S1 Fig.**
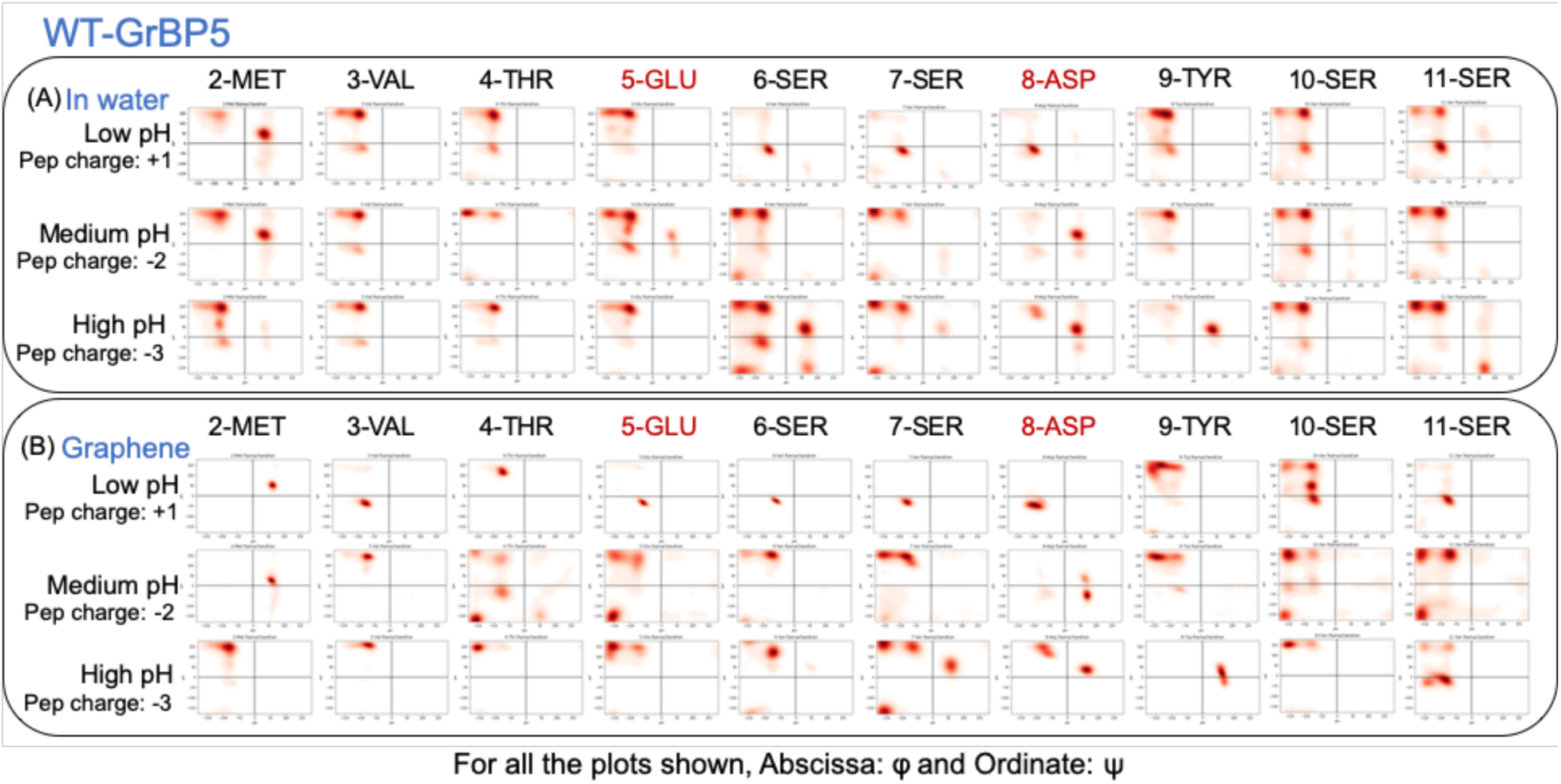
Ramachandran Plots of the interior phi-psi angle pairs of WT-GrBP5. Ramachandran plots of backbone phi-psi pairs of the interior amino acids of WT-GrBP5 from molecular dynamics simulations under low (pH < 2.24), medium (4.15 < pH < 9.6), and high (9.6 < pH < 10.10) pH conditions (A) in water; and (B) on graphene.

**S2 Fig.**
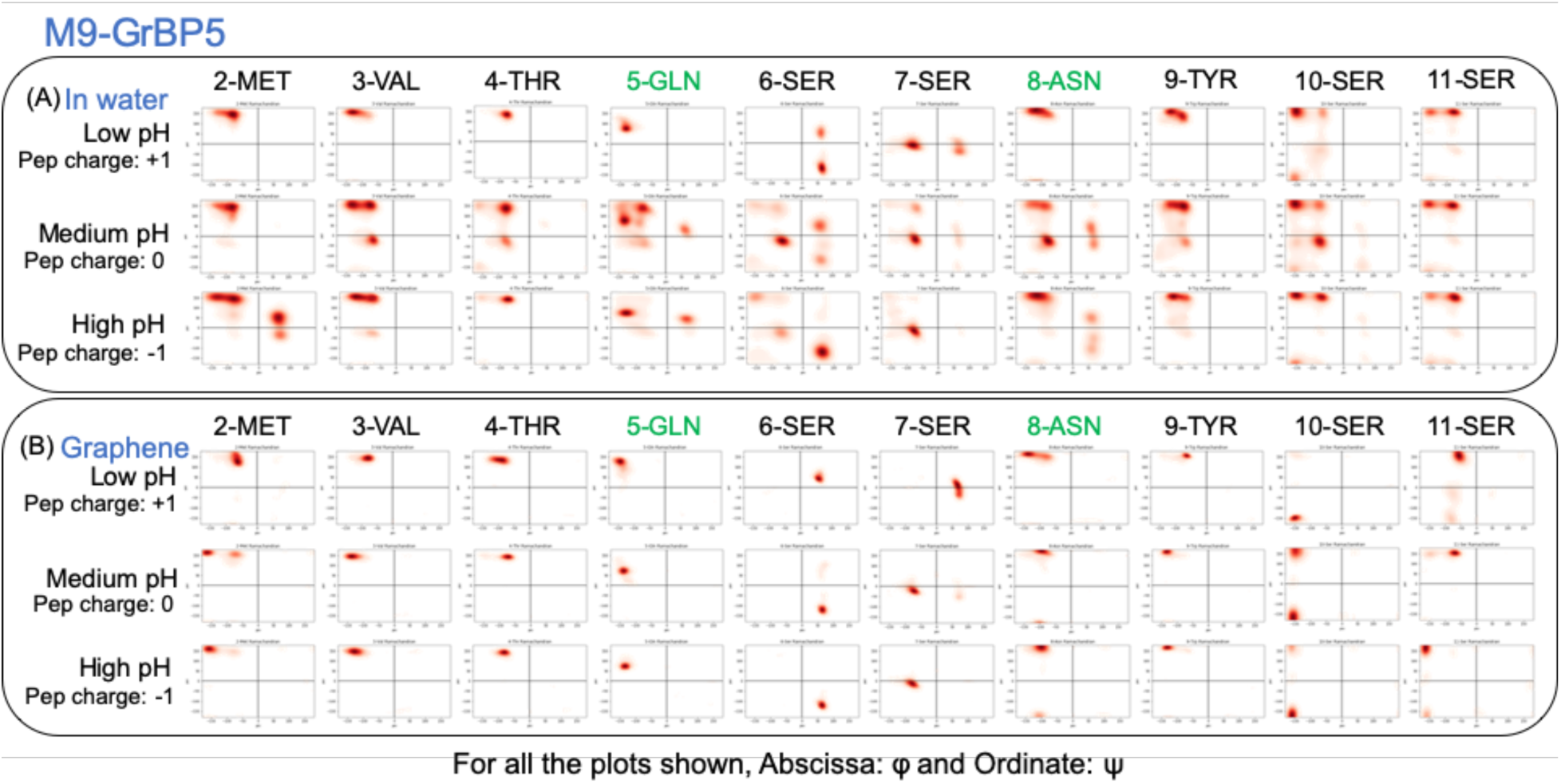
Ramachandran Plots of the interior phi-psi angle pairs of M9 GrBP5. Ramachandran plots of backbone phi-psi pairs of the interior amino acids of the charge-neutral mutant, M9-GrBP5, from molecular dynamics simulations under low (pH < 2.24), medium (2.24 < pH < 9.6), and high (9.6 < pH < 10.10) pH conditions (A) in water; and (B) on graphene.

**S3 Fig.**
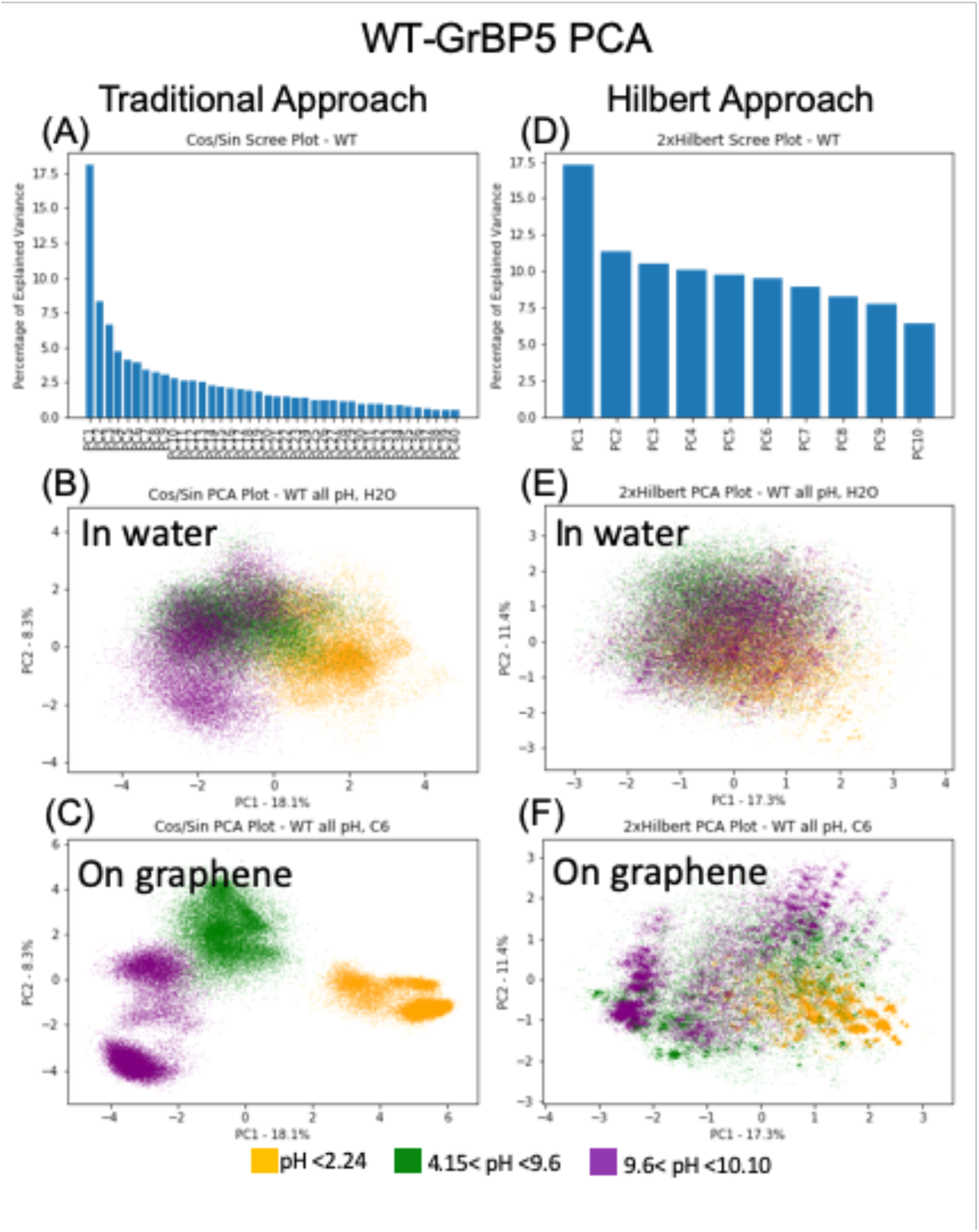
Principal Component Analysis using the Traditional Approach vs Hilbert Space-Filling Curve Approach of WT-GrBP5 in water and on graphene. (A) Scree plot from PCA of the combined linearized phi/psi data of WT-GrBP5 in water and on graphene under low (pH < 2.24), medium (4.15 < pH < 9.6), and high (9.6 < pH < 10.10) pH; (B)-(C) Scatter plot from Traditional Approach PCA, in PC1-PC2 space, of WT-GrBP5 data of the water simulations and graphene simulations, respectively, under each of the different pH conditions; (D) Scree plot from PCA of the Hilbert coordinates of the combined linearized phi/psi data of WT-GrBP5 in water and on graphene under low, medium, and high pH; (E) Scatter plot from Hilbert Approach PCA, in PC1-PC2 space, of WT-GrBP5 data of the water simulations and graphene simulations, respectively, under each of the different pH conditions. Please note WT-GrBP5 water and graphene simulations are combined into the same PCA space and are separated into two plots (for each approach) to look at them individually.

**S4 Fig.**
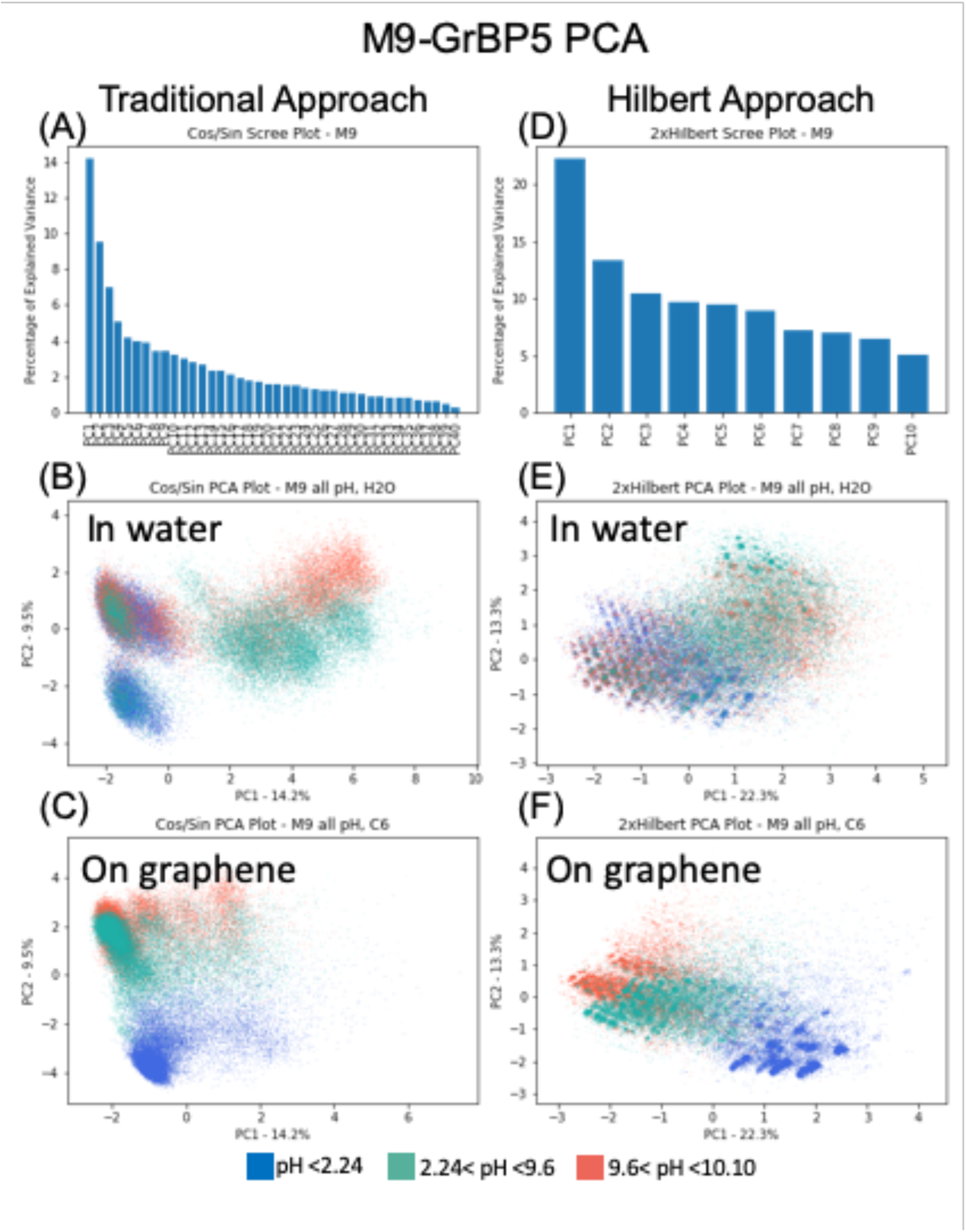
Principal Component Analysis using the Traditional Approach vs Hilbert Space-Filling Curve Approach of M9-GrBP5 in water and on graphene. (A) Scree plot from PCA of the combined linearized phi/psi data of M9-GrBP5 in water and on graphene under low (pH < 2.24), medium (2.24 < pH < 9.6), and high (9.6 < pH < 10.10) pH; (B)-(C) Scatter plot from Traditional Approach PCA, in PC1-PC2 space, of M9-GrBP5 data of the water simulations and graphene simulations, respectively, under each of the different pH conditions; (D) Scree plot from PCA of the Hilbert coordinates of the combined linearized phi/psi data of M9-GrBP5 in water and on graphene under low, medium, and high pH; (E) Scatter plot from Hilbert Approach PCA, in PC1-PC2 space, of M9-GrBP5 data of the water simulations and graphene simulations, respectively, under each of the different pH conditions. Please note that M9-GrBP5 water and graphene simulations are combined into the same PCA space and are separated into two plots (for each approach) to examine them individually.

**S5 Fig.**
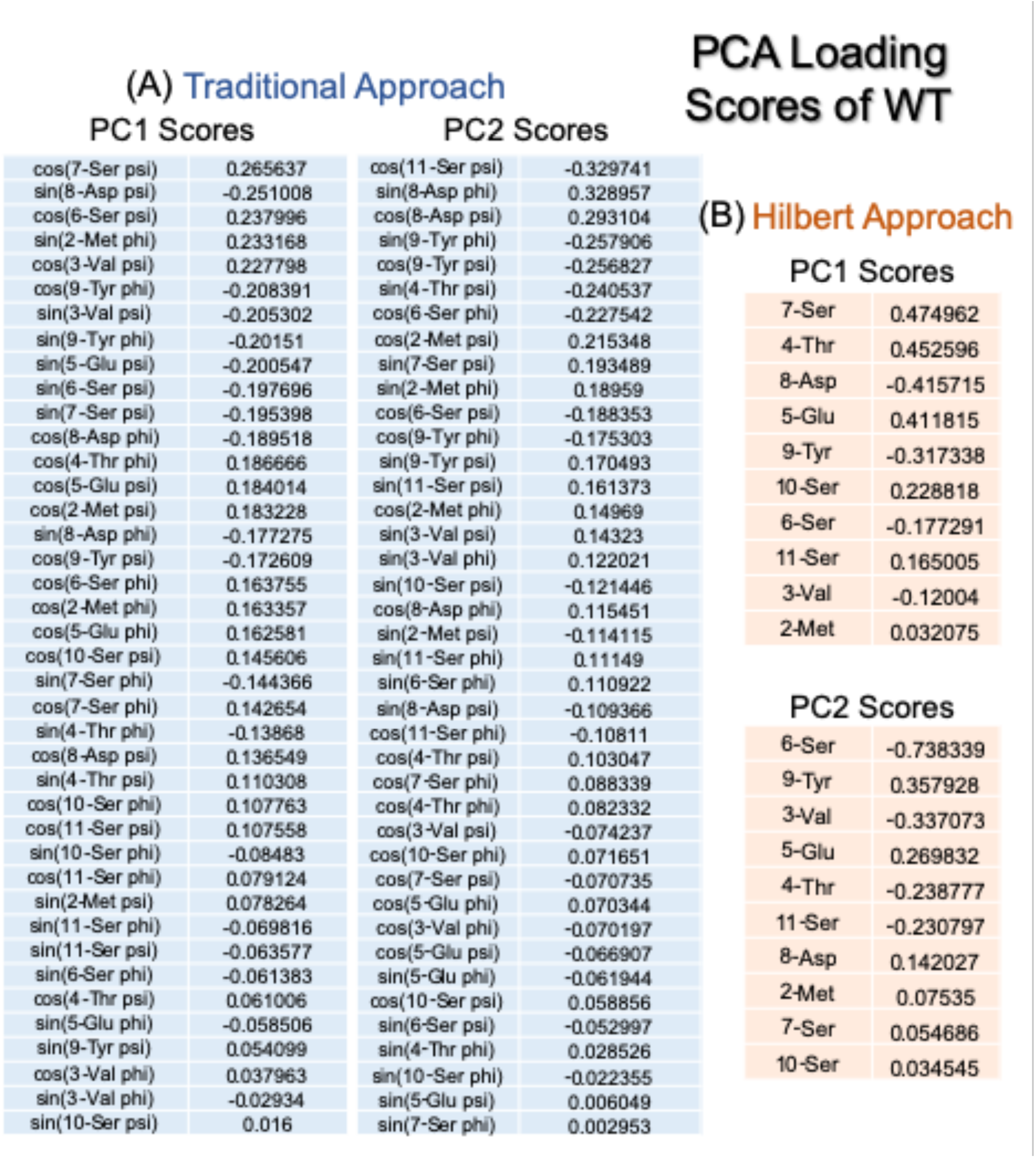
Loading Scores of WT-GrBP5 using the Traditional Approach vs Hilbert Approach. (A) Loading Scores for PC1 and PC2 from Traditional Approach PCA, using combined linearized phi/psi data from water and graphene simulations of WT-GrBP5; and (B) Loading Scores for PC1 and PC2 from Hilbert Approach PCA, using combined Hilbert coordinate phi/psi data from water and graphene simulations of WT-GrBP5.

**S6 Fig.**
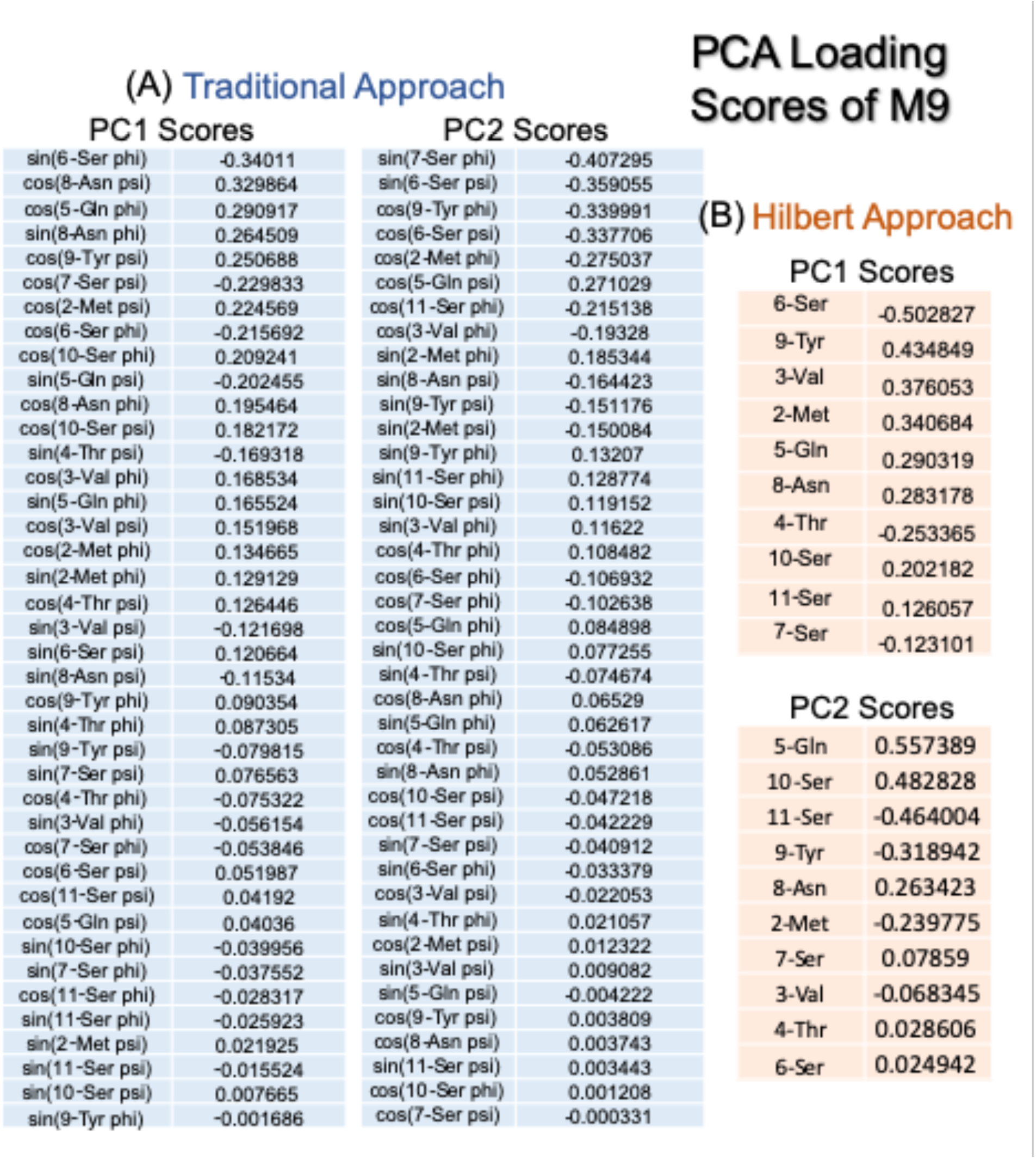
Loading Scores of M9-GrBP5 using the Traditional Approach vs Hilbert Approach. (A) Loading Scores for PC1 and PC2 from Traditional Approach PCA, using combined linearized phi/psi data from water and graphene simulations of M9-GrBP5; and (B) Loading Scores for PC1 and PC2 from Hilbert Approach PCA, using combined Hilbert coordinate phi/psi data from water and graphene simulations of M9-GrBP5.

**S7 Fig.**
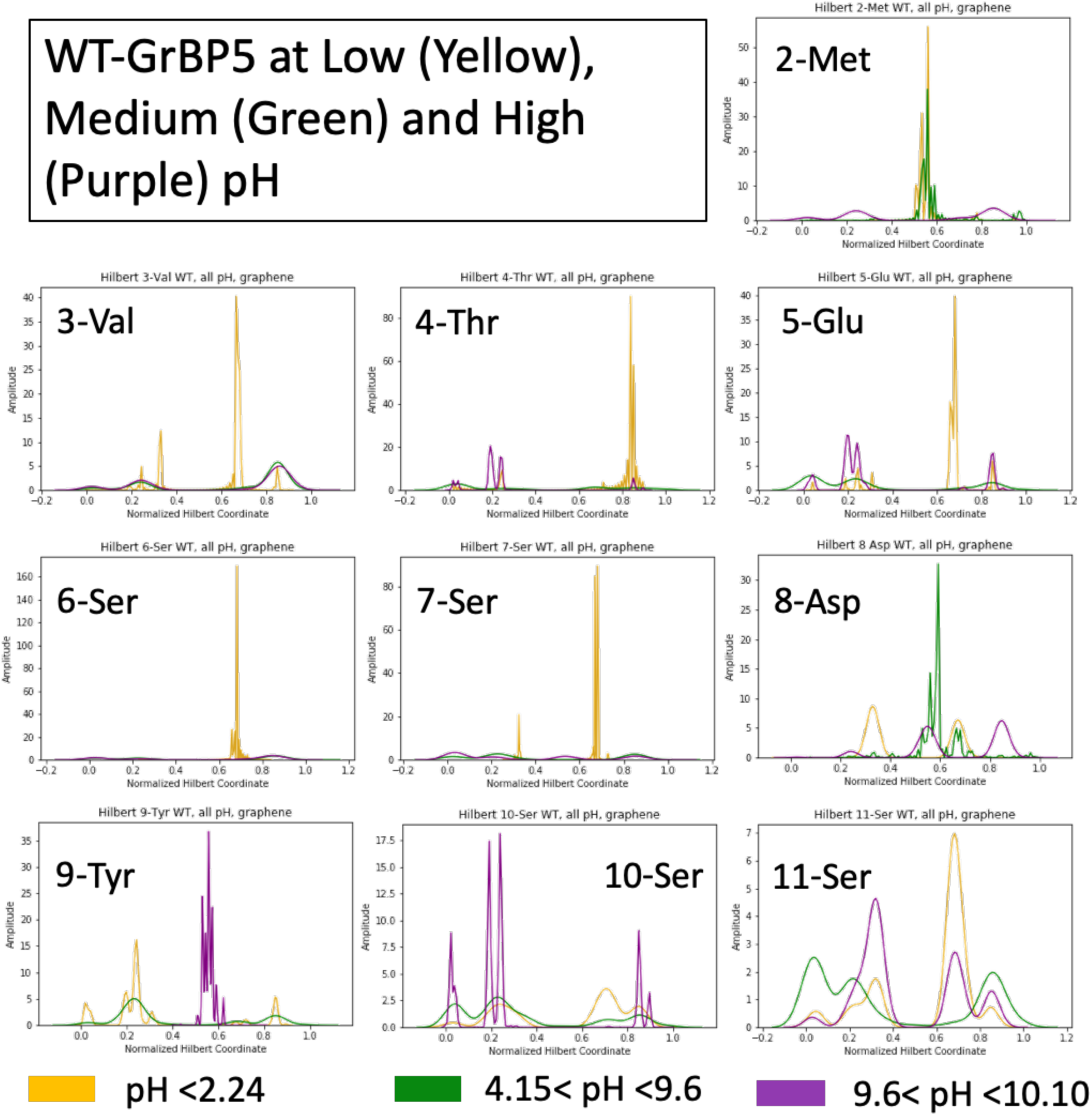
Normalized HSFC Plots of the interior phi-psi angle pairs of WT-GrBP5. Normalized HSFC distributions of backbone phi-psi pairs (consolidated into the x-axis) of the interior amino acids of WT-GrBP5 from molecular dynamics simulations under low (pH < 2.24), medium (4.15 < pH < 9.6), and high (9.6 < pH < 10.10) pH conditions on graphene.

**S8 Fig.**
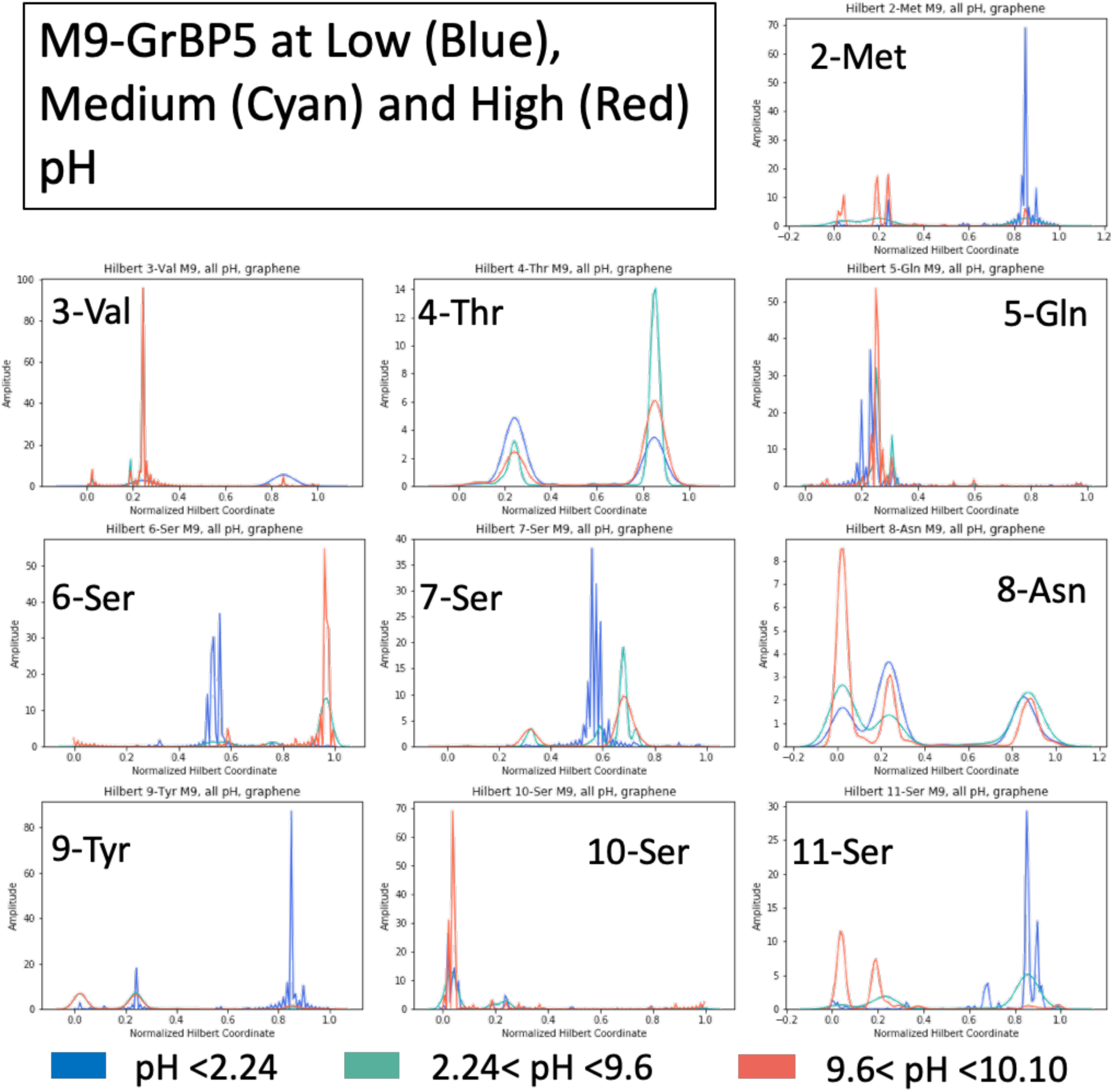
Normalized HSFC Plots of the interior phi-psi angle pairs of M9-GrBP5. Normalized HSFC distributions of backbone phi-psi pairs (consolidated into the x-axis) of the interior amino acids of M9-GrBP5 from molecular dynamics simulations under low (pH < 2.24), medium (2.24 < pH < 9.6), and high (9.6 < pH < 10.10) pH conditions on graphene.

